# A dynamic HIF1α- PPARγ circuit controls a paradoxical adipocyte regulatory landscape

**DOI:** 10.1101/2022.05.17.492374

**Authors:** Takamasa Kudo, Michael L. Zhao, Kyle Kovary, Edward L. LaGory, Markus W. Covert, Mary N. Teruel

**Affiliations:** Department of Chemical and Systems Biology, Stanford University, Stanford, California, 94305, USA; Department of Radiation Oncology, Stanford University, Stanford, California, 94305, USA; Department of Bioengineering, Stanford University, Stanford, California, 94305, USA; Department of Biochemistry and the Drukier Institute of Children’s Health, Weill-Cornell Medicine, Cornell University, New York, NY, 10065, USA

## Abstract

Hypoxia-induced upregulation of HIF1α triggers adipose tissue dysfunction and insulin resistance in obese patients. HIF1α closely interacts with PPARγ, the master regulator of adipocyte differentiation and lipid accumulation, but there are conflicting results how this co-regulation controls the excessive lipid accumulation that drives adipocyte dysfunction. Using single-cell imaging and modeling, we find that, surprisingly, HIF1α both promotes and represses lipid accumulation during adipogenesis. We show that the opposing roles of HIF1α are isolated from each other and depend on when HIF1α increases relative to the positive-feedback mediated upregulation of PPARγ that drives adipocyte differentiation. A theoretical model incorporating our findings resolves conflicting prior results and suggests that three network nodes before and after the isolation step have to be synergistically targeted in therapeutic strategies to revert hypoxia-mediated adipose tissue dysfunction in obesity.

**Teaser:** A systems biology approach detangles the effect of hypoxic and adipogenic regulators on lipid accumulation in adipocytes.

## Introduction

Obesity leads to the development of hypoxia, which is caused by increased oxygen consumption and insufficient vascularization to keep up with the expanding tissue mass (Trayhurn, 2013; Trayhurn and Wood, 2004). In response to hypoxia, cells upregulate the transcription factor HIF1α, which promotes glycolysis, vascularization, and other adaptive responses that are beneficial in allowing cells to cope with transient low oxygen conditions. However, in a state of chronic hypoxia, HIF1α signaling can become pathological, inducing inflammation and tissue fibrosis (Halberg et al., 2009; Lee et al., 2014; Seo et al., 2019).

Pathological HIF1α signaling has been shown to be a primary driver of adipocyte dysfunction and insulin resistance in obesity (Trayhurn et al., 2008; Zhang et al., 2010). Thus, understanding how to revert pathological HIF1α signaling in adipocytes could have significant impacts on treating the harmful metabolic effects of obesity. However, a paradoxical body of work reporting contradictory effects of hypoxia in adipose tissue poses a significant challenge towards developing a unified model explaining HIF1α’s role in fat cell differentiation and growth that could inform therapeutic interventions (Ban et al., 2014; Gonzalez et al., 2019; Hammarstedt et al., 2018; Lin and Yun, 2015). For example, loss-of-function experiments in mice using knockout models or dominant-negative mutants demonstrated that HIF1α inhibition could both increase and decrease fat mass and body weight (Krishnan et al., 2012; Sun et al., 2013; Zhang et al., 2010). Gain-of-function experiments in mice produced similar paradoxical results, showing that upregulation of HIF1α activity could both increase and decrease fat mass (Matsuura et al., 2013; Michailidou et al., 2015). Finally, while early in vitro adipocyte studies showed that HIF1α activation led to a failure to upregulate adipogenic genes and accumulate intracellular lipid droplets (Kim et al., 2005; Musutova et al., 2020), a more recent study found that mild hypoxia resulted in increased lipid accumulation, and that severe hypoxia inhibited lipid accumulation (Weiszenstein et al., 2016).

Taken together, these studies left the question of HIF1α’s role in adipocytes unresolved. We considered that the paradoxical findings how HIF1α leads to adipocyte dysfunction could be due to HIF1α having multiple, potentially overlapping roles in regulating lipid accumulation and in interacting with PPARγ, the master transcription regulator of adipogenesis (Caron et al., 2017; Fujisaka et al., 2013; Gaspar and Velloso, 2018; Lee et al., 2014; Shao et al., 2021; Zhang et al., 2010). To understand these roles, we integrated single-cell measurements of HIF1α, PPARγ, and lipid accumulation during adipogenesis with computational modeling of the HIF1α-PPARγ network. We show that HIF1α switches from a negative to positive role in regulating lipid accumulation in adipocytes depending on the strength of PPARγ autoregulation. We identify three critical factors: the strength of the differentiation cue, timing of hypoxic perturbation, and the strength of HIF1α activity that, if taken into account, resolve the prior paradoxical studies. Together, our results provide a unified and systems-level model for the role of HIF1α signaling in fat tissues.

## Results

### HIF1α signaling promotes lipid production, but only at high levels of PPARγ

To better understand the role of HIF1α signaling in adipogenesis, we first tested whether HIF1α levels are regulated in a differentiation-dependent manner by using rosiglitazone to induce differentiation of an established adipocyte OP9 cell line (Wolins et al., 2006). We used fluorescence microscopy to visualize the levels of HIF1α, lipid accumulation, and PPARγ, the master transcriptional regulator of adipogenesis (Lefterova et al., 2014), after four days when most cells were fully differentiated (Figure 1A, see also Materials and Methods). In agreement with previous studies (Floyd et al., 2007; He et al., 2011), HIF1α levels were strongly upregulated in response to adipogenic stimulation (Figure 1A, B). We further confirmed that HIF1α upregulation during adipogenesis is mediated by PPARγ by demonstrating that siRNA-mediated depletion of PPARγ significantly dampens HIF1α expression (Figure 1C).

**Figure 1.**
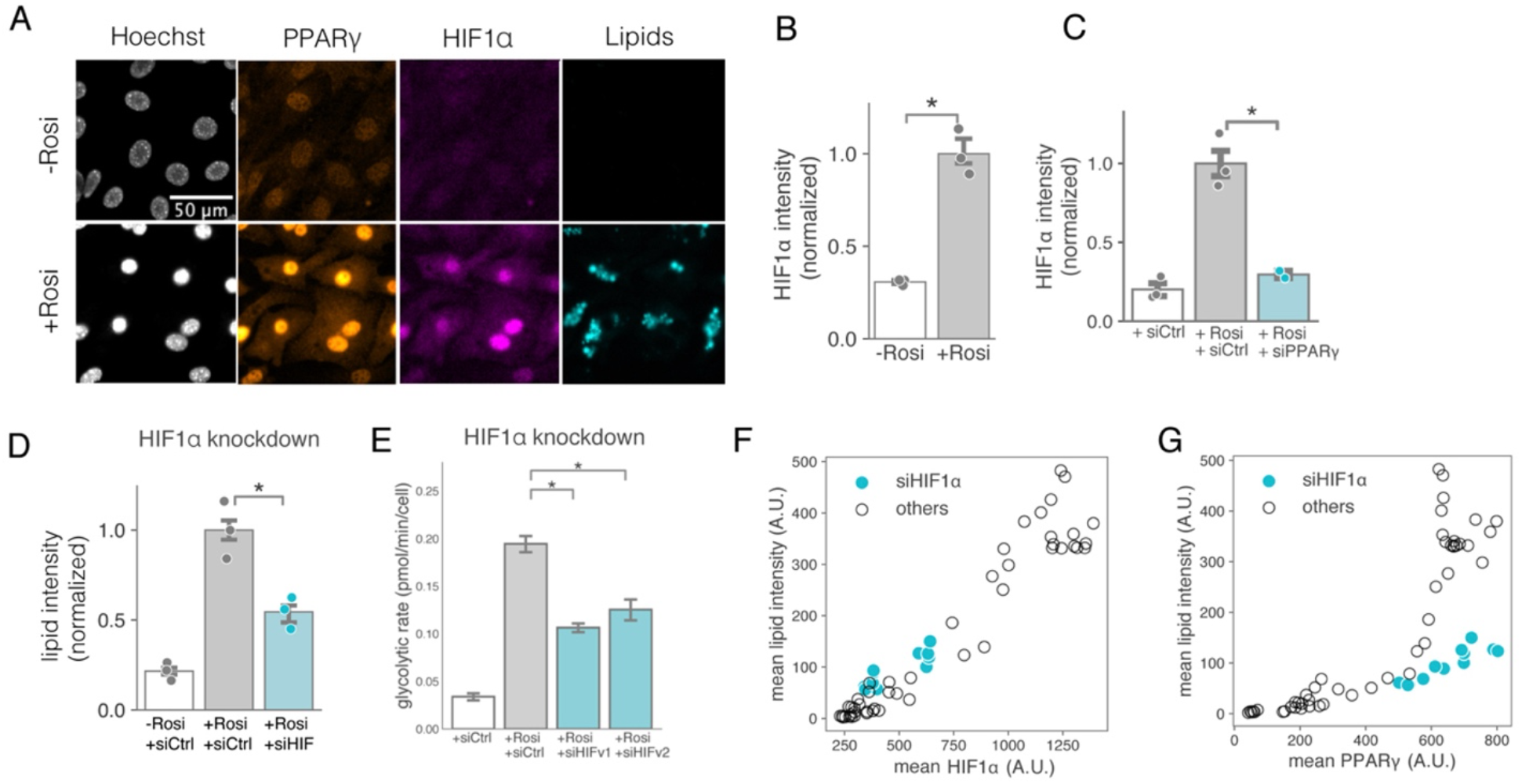
HIF1*α* signaling promotes lipid production as a downstream metabolic effector of PPARγ. (A) Representative immunofluorescence images of OP9 cells. Cells were differentiated with the 0 *µ*M (top) or 0.25 *µ*M (bottom) rosiglitazone protocol for 4 days and fixed for staining with Hoechst to visualize nuclei (gray), LipidTOX to visualize lipids (cyan), and antibodies against PPARγ (orange) and HIF1α (magenta). (B) HIF1α expression was induced upon rosiglitazone treatment. The population mean of the HIF1α intensity was quantified from immunofluorescence images. The results are normalized to their maximum value. Each dot represents a biological replicate and the bar plot is presented as means ± SD, two-tailed t test (* P < 0.05). (C) PPARγ knockdown significantly reduced the level of HIF1α expression four days after rosiglitazone stimulated differentiation. PPARγ siRNA was transfected one day before differentiation induction and HIF1α levels were assessed by immunofluorescence staining at the end of the four-day differentiation protocol. (D) HIF1α knockdown reduced the lipid level. The population mean of the lipid intensity was quantified from LipidTOX stained images. (E) HIF1α knockdown significantly reduced the rate of glycolytic flux. The flux was measured using the Seahorse Glycolytic Rate Assay. Cells were differentiated for four days (1 *µ*M rosiglitazone) with or without HIF1α siRNA transfection. siHIFv1 and siHIFv2 correspond to two individual HIF1α siRNA. Line and bar plots are presented as mean ± SD from four replicates. (F) HIF1α level showed a strong correlation with lipid level under various conditions. Cells were differentiated by either rosiglitazone or DMI differentiation protocol with siRNA targeting one of PPARγ, CEBPB, CEBPA, FABP4, HIF1α or non-targeting siRNA control. (G) HIF1α knockdown decoupled the level of PPARγ from the lipid level. (F-G) Each circle represents the mean of several hundred individual cells from a different experimental condition.

We next probed whether HIF1α plays a role in lipid accumulation during adipogenesis. Knockdown of HIF1α using siRNA led to a significant reduction of intracellular lipid levels (Figure 1D), suggesting a positive role of HIF1α in lipid synthesis. Metabolic flux assays indicated that this effect was likely mediated through an increase in glycolytic flux regulated by HIF1α (Figure 1E, S1A). To determine how robust the relationship between HIF1α and lipid synthesis is in adipogenesis, we performed a series of perturbations including siRNA knockdown of five known adipogenic genes (PPARγ, CEBPB, CEBPA, FABP4 and HIF1α) and stimulation with two differentiation signals (rosiglitazone and DMI), and measured HIF1α and PPARγ expression together with lipid accumulation for each condition. Indeed, HIF1α and lipid accumulation were positively correlated for all conditions tested (Figure 1F). Furthermore, knockdown of HIF1α resulted in a decoupling of lipid accumulation and PPARγ expression for high levels of PPARγ (Figure 1G), supporting that HIF1α regulates lipid accumulation independently of PPARγ levels in mature adipocytes.

### High levels of HIF1α can paradoxically both promote and inhibit lipid accumulation in cells in the same population

Given the positive correlation between HIF1α expression and lipid levels (Figure 1F), we next tested whether HIF1α expression is causative for lipid accumulation in differentiating preadipocytes. We generated a cell line that constitutively expresses a stable mutant of HIF1α (stblHIF), which lacks the hydroxylation sites targeted for degradation (OP9^HIF^) (Pawlus et al., 2014; Warnecke et al., 2004). To our surprise, HIF1α overexpression significantly reduced the population-averaged lipid levels to a similar extent as achieved by HIF1α knockdown (Figure 2A and 1D). To confirm this observation, we used a well-known chemical inhibitor (PX-478) and activator (FG-4592) of HIF1α and found that both inhibition and activation of HIF1α reduced lipid accumulation (Figure 2B). Thus, HIF1α can have opposite signaling roles in the regulation of lipid accumulation reminiscent of the results reported in the mouse studies. In this way, our results argue that the experimental system we developed can be used to dissect the mechanism underlying the paradoxical experimental observations in the literature which we described above.

**Figure 2.**
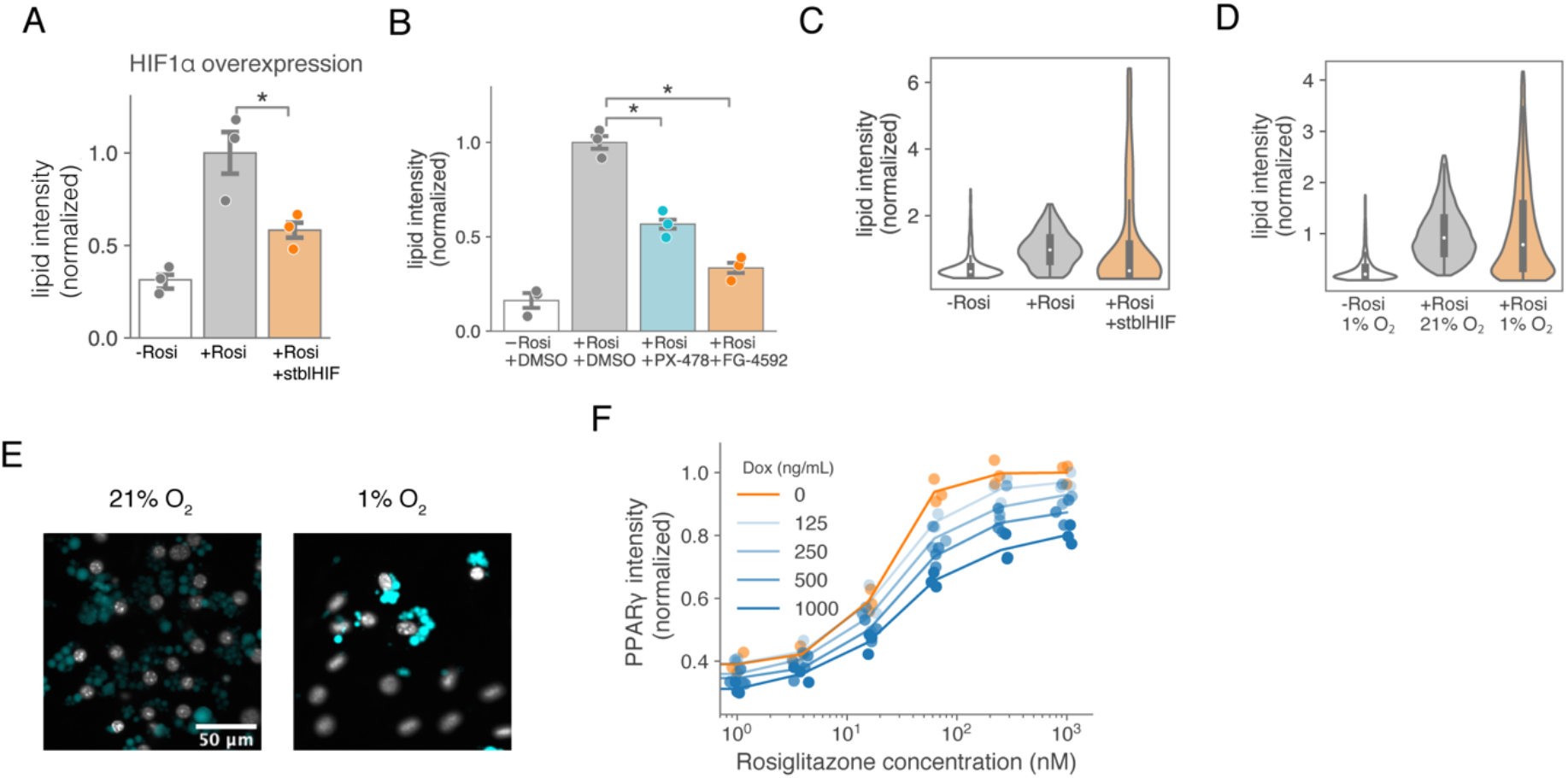
High levels of HIF1*α* can both paradoxically promote or inhibit lipid accumulation in cells in the same population. (A-E) OP9 preadipocyte cells were differentiated by addition of rosiglitazone and then fixed and stained at Day 4 when cells were fully differentiated. LipidTOX was used to quantify lipids. (A) HIF1α was overexpressed during adipogenesis by adding doxycycline to stblHIF cells together with the rosiglitazone differentiation stimulus. Cells showed reduced lipid accumulation at Day 4. (B) Wildtype OP9 preadipocyte cells were differentiated and treated with 50 *µ*M PX-478 or 50 *µ*M FG-4592 for HIF1α inhibition and activation, respectively. Both chemical inhibition and activation of HIF1α resulted in reduced lipid levels at Day 4. (C-D) The normalized lipid intensity in each of the thousands of individual cells per condition were plotted in violin plots. The plotted shape width represents the probability density of the data at the different lipid intensity values. (C) HIF1α overexpression in cells differentiated with 0.25 *µ*M rosiglitazone resulted in a long-tail distribution of lipid levels. (D) Cells differentiated with 1 *µ*M rosiglitazone under hypoxic conditions also showed a long-tailed distribution of lipid levels. Cells were differentiated for 6 days either in 21% O2 (gray, normoxia) or 1% O2 (orange, hypoxia). (E) Representative images of cells differentiated under hypoxic conditions in (B). Nucleus marked by Hoechst (gray) and lipids marked by LipidTOX Deep Red (cyan). (F) HIF1α overexpression reduced PPARγ level in a doxycycline concentration-dependent manner. Each dot represents one replicate averaging several hundred individual cells, and the line plot shows the mean of replicates.

When using violin plots to examine how much lipid each individual cell accumulated during differentiation (Figure 2C), we noticed a long-tail distribution of lipid intensities when HIF1α was overexpressed, meaning that a small subset of cells had greatly increased lipid levels rather than the low, HIF1α-inhibited lipid levels seen in the other cells (Figure 2C). We further verified the long tail distribution by differentiating cells under hypoxic conditions (Figure 2D and E). The long-tailed distribution suggested that HIF1α may play a dual role in adipogenesis, in which HIF1α expression inhibits lipid accumulation in most cells but enhances lipid accumulation in a small subpopulation of cells.

As our previous results had established a strong positive role of HIF1α in lipid accumulation (Figure 1), we next examined its negative role. We considered that HIF1α may downregulate PPARγ as reported in some studies (Krishnan et al., 2012; Shao et al., 2021; Yun et al., 2002) which then may indirectly reduce lipid accumulation. We modified the stblHIF construct such that its expression can be induced by doxycycline (OP9^treHIF^). We used doxycycline and control conditions to compare the effect of high and low HIF1α before treating cells with rosiglitazone for 24 hours to induce PPARγ expression. This analysis showed that increased HIF1α expression can inhibit PPARγ expression induced by the adipogenic stimulus (Figure 2F).

### HIF1α has two roles during adipogenesis that can be temporally separated

Given the opposing roles of HIF1α in regulating lipid accumulation, we hypothesized that the timing of signaling during adipogenesis might be critical. Therefore, we measured levels of HIF1α, PPARγ, and lipids each day during the time course of differentiation. We found that the level of HIF1α was low early during differentiation but sharply increased after day 2 (Figure 3A, see also S2A and S2B for controls with or without insulin). To test the specific impact of HIF1α expression timing on lipid accumulation, we then treated OP9^treHIF^ cells with doxycycline at different times relative to the differentiation stimulus (Figure 3B). Markedly, induction of HIF1α prior to adding rosiglitazone inhibited lipid accumulation, while induction of HIF1α at subsequent time points increased lipid accumulation. We made similar observations when exposing wildtype OP9 cells to hypoxia on different days using a similar protocol (Figure 3C). These results showed that early in the differentiation process, HIF1α inhibits lipid accumulation. However, later in the differentiation process, HIF1α increases lipid accumulation.

**Figure 3.**
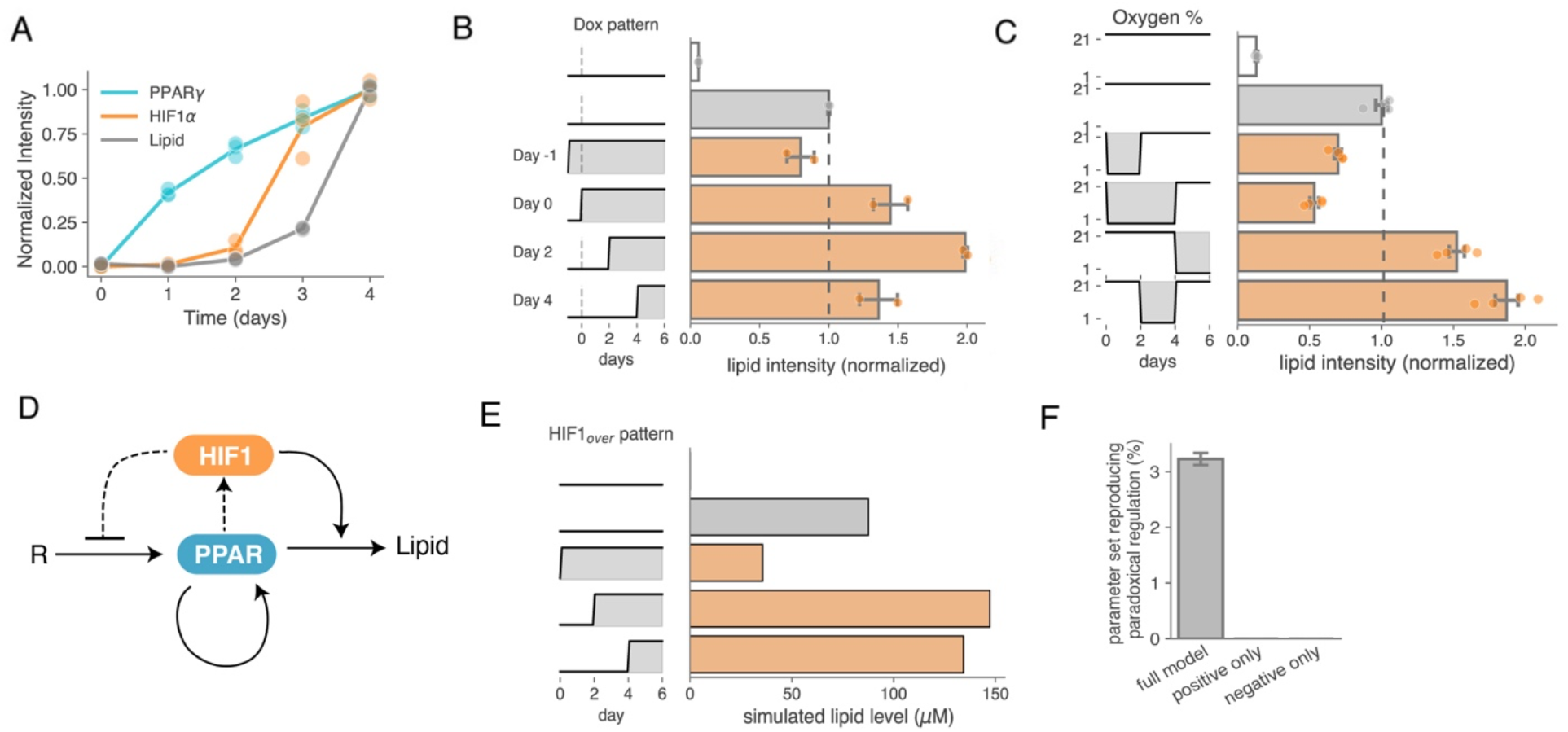
HIF1*α* has two roles during adipogenesis that can be temporally separated. (A) OP9 cells were induced to differentiate by adding 100 nM rosiglitazone and then fixed and immunocytochemistry was carried out. PPARγ levels increased immediately, but HIF1α increased acutely only two days later. Each dot represents the mean of several hundred individual cells. Results are normalized to the non-perturbed condition. (B) Induction of HIF1α expression early in adipogenesis reduced the lipid level, while the late overexpression increased the lipid level measured at day 6. OP9^treHIF^ cells were differentiated with rosiglitazone (100 nM for grey bar, 0 nM for white bar) and fixed at day 6 for staining. stblHIF expression was induced by 0.4 *µ*g/ml doxycycline for different timing and duration. (C) Early exposure to hypoxia (1% O2) reduced the lipid level, while the late exposure increased the lipid level measured at day 6. OP9 cells were differentiated with rosiglitazone (0.25 *µ*M for grey bar, 0 nM for white bar) and fixed at day 6. Cells were exposed to 1% O2 for different timing and duration where the shaded region represents hypoxia. (B-C) Results are normalized to the non-perturbed condition, which is also indicated as a dashed line. (D) Scheme of the mathematical model. (E) The simulation reproduced the timing dependency of the system. Mean lipid levels were calculated from 5000 cells differentiated at 0.25 *µ*M R for six days. HIF1α overexpression (HIF_over_=1) was applied at 0, 48, or 96 h (shaded pattern on the left panels). (F) Through the randomized parameter search, the fraction of parameter sets that could recapitulate the timing-dependent paradoxical regulation was obtained for each model structure. Multiple simulation runs (30,000 searches for four times) are presented as mean ± SD. The models lacking either the positive or negative regulation from HIF1α were unable to reproduce the paradoxical observed experimentally in (B) and (C).

Previous work demonstrated that PPARγ levels irreversibly switch from low to high when cells commit to differentiate into adipocytes (Bahrami-Nejad et al., 2018; Zhao et al., 2020). This differentiation commitment point precedes most lipid accumulation and is critically dependent on multiple positive feedback loops centered on PPARγ (Ahrends et al., 2014; Bahrami-Nejad et al., 2018). We wondered whether and how the regulation of lipid accumulation by HIF1α during adipogenesis depended on the state of this PPARγ autoregulation circuit. To explore this, we constructed a computational model that integrates PPARγ autoregulation with the paradoxical dual regulation of HIF1α (Figure 3D; supplementary text). In this model, the rate of lipid accumulation is described as a function of both HIF1α and PPARγ (Figure 1F and G), while HIF1α also inhibits the rate of PPARγ synthesis (Figure 2 and 3A-C)(Shao et al., 2021; Yun et al., 2002). PPARγ activation is initiated by a differentiation stimulus (R) (rosiglitazone in our experimental system). We incorporated previously published terms and parameter values as available (Table S1) and fitted the remaining parameters. To fit parameters, we relied on our experimental data which showed that early HIF1α overexpression reduces adipocyte lipid levels while later HIF1α overexpression increases them. Using the experimental findings as selection criteria, and by including known steady-state values of HIF1α and PPARγ (Table S1), we derived parameter values (supplementary text) that recapitulated (i) how changes in HIF1α expression at different times during differentiation regulate lipid accumulation in opposite directions (Figure 3E), and (ii) that PPARγ is autoregulated in a bistable manner to trigger differentiation (Figure S3A-E).

We compared our model in Figure 3D with two additional models that lacked either (1) the HIF1α-mediated inhibition of PPARγ synthesis or (2) the induction of lipid synthesis by HIF1α. Indeed, these models lacking either the negative or the positive HIF1α regulation of lipid accumulation could not recapitulate the timing-dependent changes (Figure 3F). To test whether the opposing roles of HIF1α before and after PPARγ autoregulation are required to explain the paradoxical effect of HIF1α on lipid accumulation,

### Positive feedback centered on PPARγ functionally isolates the inhibitory and activating roles of HIF1α

We next wanted to understand what might be responsible for temporally separating the two opposing roles of HIF1α in lipid accumulation during adipogenesis. Because we had found that elevated PPARγ is needed for HIF1α to induce lipid accumulation late in adipogenesis (Figures 1G and 3A), we wanted to test if the temporal separation is a result of HIF1α having an opposing earlier role by preventing the positive-feedback-mediated upregulation of PPARγ. Thus, we next used the model from Figure 3D to explore how HIF1α influences the upregulation of PPARγ when PPARγ levels are initially low. To do so, we used a sensitivity analysis in which we introduced mild perturbations to test for the connection between HIF1α upregulation and positive-feedback regulation of PPARγ. Specifically, we simulated lipid accumulation in response to a differentiation stimulus (1) when HIF1α is partially activated early in differentiation, (2) when the PPARγ positive feedback is attenuated, or (3) when both perturbations are combined (Figure 4A). We chose to use these perturbations since they can be used experimentally to test for whether HIF1α can indeed suppress the non-linear positive feedback early in differentiation to influence lipid accumulation.

**Figure 4.**
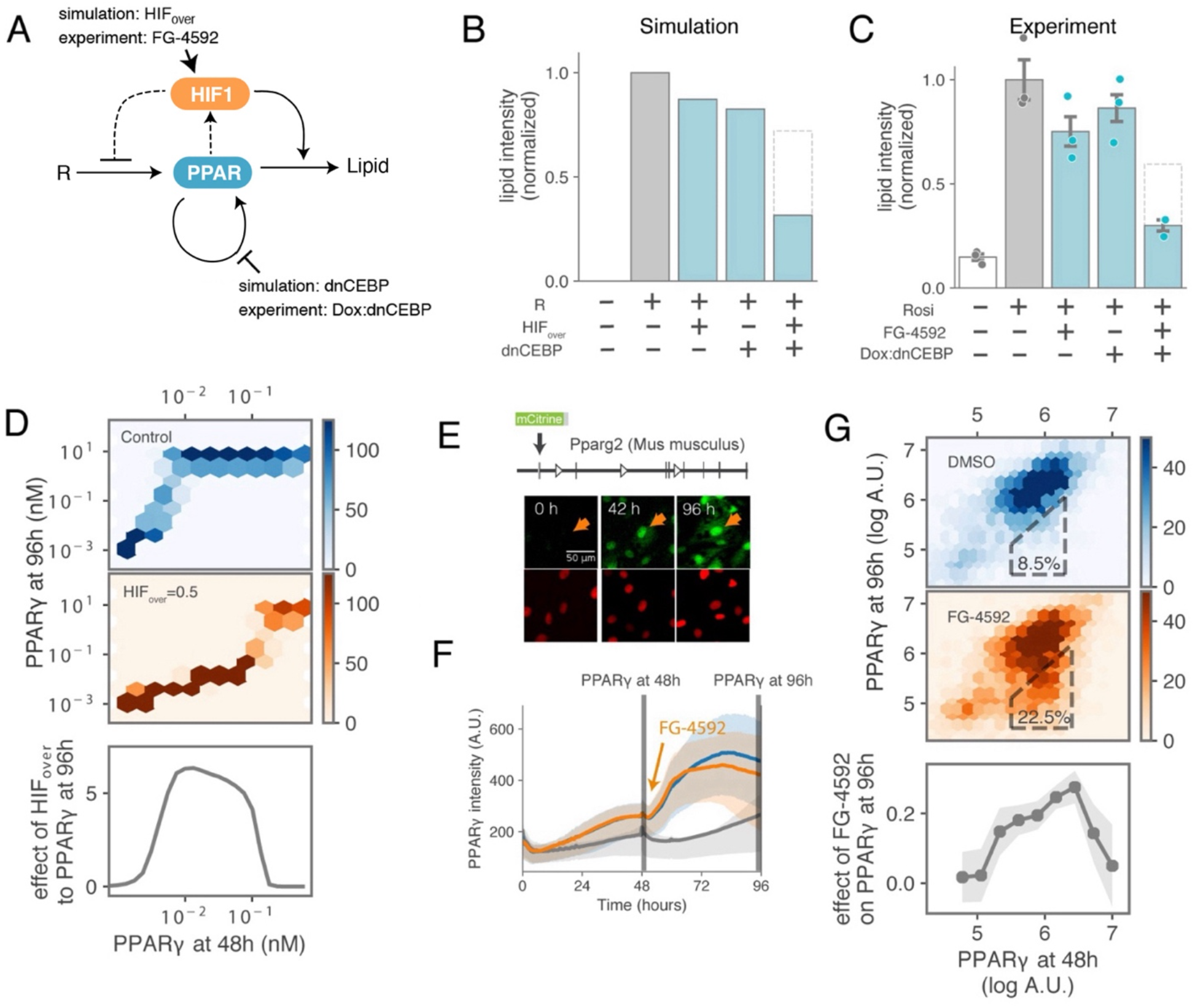
PPARγ positive feedback functionally isolates the inhibitory and activating roles of HIF1*α*. (A) Cartoon of the perturbation scheme used to compromise the PPARγ positive feedback and induce HIF1α activation. (B) The model simulation showing the feedback compromisation by dnCEBP substantiated the negative effect from HIF1α overexpression. The presence of R, HIFover, dnCEBP indicates R=0.25, HIFover =0.5, Ci =0.33 in the model. The box with a dashed line assumed the linear effect of HIF1α overexpression and dnCEBP expression. (C) The experimental results reproduced the model simulation. Cells were differentiated (1 *µ*M rosiglitazone) and treated with either 12.5 *µ*M FG-4592 or 0.4 *µ*g/ml doxycycline, or both. Each dot represents a biological replicate and the bar plot is presented as means ± SD. (D) Computationally-simulated relationship between PPARγ at 48h and 96 h predicts that HIF1α overexpression affects only cells with intermediate PPARγ before overexpression. Hexagons are colored according to cell density. Darker colors of hexagons mark more cells with the respective 48h and 96h PPARγ levels. The difference in mean PPARγ level at 96h between control (blue) and HIF1α overexpression (orange) conditions is plotted (bottom). (E) Schematics of the live-cell experiment. PPARγ-mCitrine (green) and H2B-Turquoise (red) fluorescence intensities were imaged for 96 h after adding rosiglitazone to induce differentiation. The orange arrow indicates a representative cell tracked over time. (F) FG-4592 treatment (25 *µ*M) at 48 h after differentiation mildly inhibited PPAR accumulation. The line represents population means and the shade represents values between the 25th and 75th percentile. Cells were treated with rosiglitazone (0.25 *µ*M) (blue), rosiglitazone and FG-4592 (orange) or without rosiglitazone (gray). (G) Experimentally-measured relationship between PPARγ at 48 h and 96 h. Cells were treated with DMSO (control, blue) or FG-4592 (orange) at 48 h. Hexagons are colored according to cell density. Box outlines highlight cells with intermediate PPARγ levels that are most affected by FG-4592. The difference in mean PPARγ level at 96 h between DMSO and FG-4592-treated conditions is plotted (bottom). The shaded area represents standard error of the mean estimated from 100 bootstrapping steps, sampling a half population each time.

The model simulations predicted that lipid accumulation is only mildly impacted by either perturbation alone but is severely suppressed when both perturbations are applied simultaneously (Figure 4B). To experimentally validate the model predictions, we developed a system in which we could both activate HIF1α using small molecules and attenuate PPARγ autoregulation by manipulating expression of the C/CAAT Enhancer binding proteins, CEBPα and CEBPβ, which have been previously shown to be critical PPARγ feedback regulators. To target the CEBP feedback loop, we generated a stable OP9 preadipocyte cell line in which we could induce a dominant-negative form of CEBP (dnCEBP) with doxycycline (Musunuru et al., 2010; Olive et al., 1996; Zhang et al., 2004). We added rosiglitazone to the cells to induce adipogenesis together with (1) FG-4592 to activate HIF1α, (2) doxycycline to induce expression of dnCEBP and attenuate PPARγ autoregulation, or (3) both perturbations together (Figure 4C). Four days later when the cells were differentiated, we measured lipid accumulation and found that neither perturbation by itself resulted in a large reduction of lipids. However, a significant synergistic decrease in lipids was observed after co-treatment with both FG-4592 and doxycycline, as predicted by the model simulations. The synergism between early HIF1α signaling and PPARγ autoregulation in regulating lipid accumulation strongly support that HIF1α has a direct role early in adipogenesis to suppress the commitment to differentiate.

Since PPARγ autoregulation triggers an irreversible commitment step in the differentiation process (Bahrami-Nejad et al., 2018), our finding of opposing roles of HIF1α before and after differentiation commitment argues that differentiatino commitment has an insulating role - namely to protect the differentiated cells from the initial HIF1α-mediated suppression of PPARγ. Insulation explains how increased HIF1α expression greatly increases lipid accumulation only after commitment (Figure 1). Such an insulating effect where signal regulation can change before and after a bistable switch has been recently reported in two cell systems that have bistable switches (Araujo et al., 2016; Atay et al., 2016).

### The insulation effect is closely associated with high PPARγ levels in heterogeneous single cells

Single-cell levels of PPARγ have been shown to greatly vary between cells in a population, which causes a great variation in the timing of differentiation commitment in differentiating cells (Ahrends et al., 2014). This heterogeneity lets us explore how cells with different levels of PPARγ respond to an increase in HIF1α expression. We simulated a dynamic time course in which cells were induced to differentiate at 0h. Forty-eight hours later, when many of the cells had increased PPARγ to levels that triggered PPARγ autoregulation (Bahrami-Nejad et al., 2018), we upregulated HIF1α in the cells and continued the simulations from 48 to 96 hours. We visualized the inhibitory effect of HIF1α as a function of PPARγ at 48 hours by computing the difference between the simulated PPARγ levels at 96 hours, with or without HIF1α overexpression (Figure 4D, top two panels).

The simulations showed that cells that already had a high PPARγ concentration at 48 hours kept their high PPARγ concentration at 96 hours even when HIF1α was overexpressed (blue versus orange in Figure 4D, comparing cells at the right top). However, in cells with intermediate PPARγ concentrations at 48 hours, HIF1α overexpression sharply suppressed the normally seen maximal PPARγ increase observed in control cells at 96 hours (difference highlighted in the bottom plot, Figure 4D). In other words, our model results predicted that the inhibitory effect of HIF1α on differentiation is biphasic: cells with high PPARγ are already differentiated and the multiple positive feedbacks to PPARγ are strongly engaged so that HIF1α can no longer suppress PPARγ levels. Furthermore, those cells with very low PPARγ at 48 hours would not differentiate anyhow during that time. Thus, the model predicted that only cells with intermediate PPARγ levels are sensitive to being suppressed by HIF1α overexpression at 48 hours.

To test this prediction experimentally, we carried out live-cell imaging to measure PPARγ levels in individual cells over time. We used a previously generated OP9 cell line expressing endogenous PPARγ fused to mCitrine(YFP) (Figure 4E) (Bahrami-Nejad et al., 2018). The cells were also stably transfected with H2B-mTurquoise(CFP) as a nuclear marker to facilitate cell tracking and analysis. We stimulated the cells with rosiglitazone to induce differentiation, and then 48 hours later, when many of the cells had increased PPARγ to levels that triggered PPARγ autoregulation, we added the HIF1α―activator, FG-4592, to the cells and imaged for another 48 hours. The time-lapse images were then analyzed to obtain single-cell time traces of PPARγ intensity (Figure 4F). To understand how HIF1α activation affected PPARγ levels in the individual cells in the population, we compared the PPARγ level of each cell at 48 hours (before FG-4592 treatment) with its PPARγ level at 96 hours (Figure 4G, top two panels). Markedly, the cell density plots showed the biphasic relationship predicted by the computational model (Figure 4D) in which activation of HIF1α strongly suppressed 96-hour PPARγ levels in cells that had an intermediate PPARγ level at 48 hours, but only minimally affected cells that had either low or high PPARγ at 48 hours (Figure 4G, box outlines and bottom panel). Thus, our simulation and experimental results show that HIF1α suppresses PPARγ only before cells commit to differentiate and PPARγ autoregulation strongly engages. When cells have intermediate or higher levels of PPARγ, HIF1α activation does not anymore affect PPARγ levels. These results support the model that PPARγ autoregulation and differentiation commitment isolate HIF1α’s early suppression of PPARγ levels from the later HIF1α action to accumulate lipid.

### Activation of HIF1α controls lipid accumulation in opposing directions depending on the strength of the differentiation stimulus

Our data thus far shows that HIF1α only suppresses PPARγ levels and commitment before cells commit but has no effect on PPARγ levels after commitment. In its second role, HIF1α is increasing lipid accumulation but only in cells with high levels of PPARγ that already committed. Since PPARγ levels are initially increased primarily by external differentiation cues (Rosen et al., 2000), it would be important to understand how the strength of the adipogenic cue synergizes with HIF1α to increase PPARγ levels and ultimately regulate lipid accumulation. To examine the synergy, we carried out simulations at different differentiation stimulus strengths (R) and HIF1α overexpression levels and calculated the mean lipid level from 5000 cells for each condition. As expected, the parameter landscape showed accumulated lipid levels that varied significantly depending on differentiation stimulus strength and HIF1α expression level (Figure 5A, left). To better visualize the relationship, we selected out of the landscape two levels of HIFα overexpression (blue and orange dotted lines) and compared them in the same plot (Figure 5A, middle). Our model predicted that HIF1α overexpression (orange trace) results in a need for much higher adipogenic stimulus R to increase the lipid level relative to the control condition (blue trace) but at the same time leads to much higher lipid levels for the high ranges of adipogenic stimuli. To experimentally test this prediction, we titrated the concentration of rosiglitazone used to differentiate OP9^treHIF^ cells in the presence or absence of doxycycline. We found a close match between the experimental and simulation results (Figure 5A, right panel), supporting that HIF1α opposes lipid accumulation for weak adipogenic stimuli but increases lipid accumulation for strong adipogenic stimuli.

**Figure 5.**
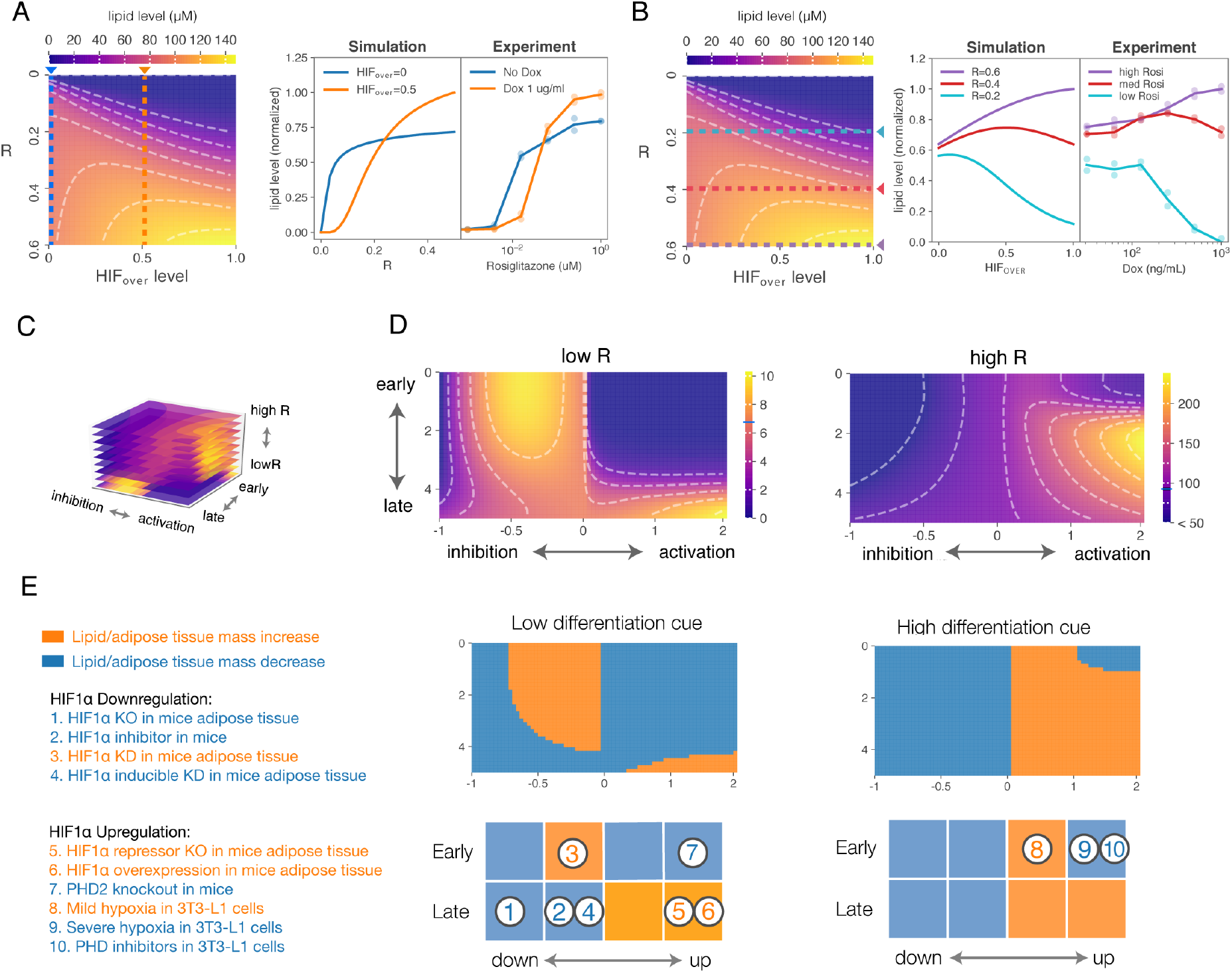
HIF1*α* perturbation revealed the paradoxical regulatory landscape of adipogenesis. (A) (left heatmap) The simulated landscape varying R and HIF1α overexpression levels. Color gradients indicate the mean lipid concentration calculated from 5,000 cells at day six. The colors of dashed lines correspond to the following panels. (right panels) The role of HIF1α overexpression is dependent on the differentiation cue. For each different level of R, the mean lipid level was simulated for the cells that were treated with (orange) or without HIF1α overexpression (blue) (left panel). The experiment reproduced the same trend (right panel), where OP9^treHIF^ cells were differentiated with different levels of rosiglitazone (0.001, 0.004, 0.015, 0.063, 0.25, 1 *µ*M), with or without HIF1α induction through doxycycline. Each experimental and simulation result was normalized to its maximum value. Each dot represents one replicate and the line plot shows the mean of replicates. (B) (left heatmap) The simulated landscape varying R and HIF1α overexpression levels. The colors of dashed lines correspond to the following panels. (right panels) The emergence of increasing, decreasing and biphasic trends by changing the differentiation cue. Mean lipid levels with varying HIFover were simulated at three different levels of R (0.2 for cyan, 0.4 for red, 0.6 for purple line). For the experimental validation, OP9^treHIF^ was differentiated by rosiglitazone (0.015 *µ*M for cyan, 0.063 *µ*M for red and 1 *µ*M for purple circles) with different concentration of doxycycline to overexpress HIF1α (0.03, 0.06, 0.13, 0.25, 0.50, 1.00 *µ*M). Each experimental and simulation result was normalized to its maximum value. (C) The 3D landscape of lipid production is dependent on the direction of HIF1α perturbation (inhibition vs activation), the timing of perturbation (early vs late), and the strength of differentiation cue (low vs high R). (D) The effect of HIF1α knockdown and overexpression at different timing on mean lipid production at two different levels of R (0.01 for left, 0.4 for the right panel). Each heatmap color is scaled within each level of R for visualization. The left half of each panel (corresponding to -1 to 0) indicates HIF_kd_ values, where -1 indicates the complete inhibition. The right half of each panel (corresponding to 0 to 2) indicates HIF_over_ values, where 1 indicates the two-fold expression relative to the steady-state HIF1α level (0.025 *µ*M) at R=1. Note that 0 on the x-axis represents endogenous HIF1 levels without any over or under expression. The blue lines on the color bars indicate the lipid level with no perturbation. Y-axis (early to late) indicates the timing of perturbation (days). (E) Perturbation landscape explains the past observations. The landscapes were first binarized to represent an increase or decrease in lipid accumulation relative to the no perturbation (top panels). Then they are reduced to the coarsely divided 2×4 grid (bottom panels: the column positions correspond to HIF_kd_=-1, HIF_kd_=-0.5, HIF_over_=1, HIF_over_=2; and the row positions correspond to day 0 and day 5 to represent the early and late timing, respectively).

We next used the same landscape to analyze the simulations in a different way to understand if and how the strength of the adipogenic stimulus matters for lipid accumulation. We examined at three different strengths of the differentiation stimulus R how increased HIF1α signaling changes lipid accumulation (purple, red, and cyan lines in Figure 5B, left). When R was high (purple), our simulations predicted that HIF1α overexpression would lead to an increase in overall lipid accumulation. When R was low (cyan), HIF1α overexpression was predicted to reduce lipid accumulation. Most interestingly, intermediate levels of R (red) produced a biphasic response, in which the amount of accumulated lipid levels increased with HIF1α overexpression up to a certain point (HIFover≃0.5), but decreased for overexpression levels beyond that point. To test these predictions, we differentiated cells with low, medium, and high concentrations of rosiglitazone and induced different degrees of HIF1α expression (Figure 5B, right). We found a remarkable agreement of the predicted biphasic behavior with weak adipogenic stimulus showing decreasing lipid accumulation with increasing HIF1α expression and strong adipogenic stimulus showing increasing lipid accumulation with increasing HIF1α expression. The medium strength of adipogenic stimulation caused for low-to-intermediate HIF1α expression an increase in lipid accumulation and for intermediate-to-high expression a decrease, again as predicted by the simulation (circles in Figure 5B).

To further test the model predictions that the strength of the differentiation stimulus synergizes with HIF1α expression to regulate lipid accumulation, we titrated the HIF1α inhibitor PX478 together with rosiglitazone and again showed that inhibition of HIF1α expression has biphasic characteristic in both the simulation and experiments (Figure S4A, B). We thus conclude that the role of HIF1α in lipid accumulation is controlled by the strength of the differentiation cue. The rationale for this synergistic regulation is that the differentiation stimulus can trigger PPARγ autoregulation and differentiation commitment which decides whether a cell becomes insulated so that HIF1α loses its ability to suppress differentiation and is left with only its role in promoting lipid accumulation.

### Biphasic response is an emergent property of a paradoxical architecture

Thus far, we learned that both the timing of overexpression and the magnitude of differentiation cue impacts the degree of PPARγ insulation, thereby affecting outcomes from HIF1α perturbation. Given this insight, we next wondered if the apparent role change of HIF1α arises as an emergent property of a paradoxical circuit. To explore this point further, we created a simplified population dynamics model that explicitly models the effect of insulation.

In this model, we consider two cell states: PPAR_low_ and PPAR_high_. PPAR_low_ cells differentiate to PPAR_high_ cells at a differentiation rate r_H_ (Figure S5A). The total amount of lipid is described as the production of PPAR_high_ cells and lipid production rate k_H_. To represent the paradoxical dual action, HIF1α overexpression (H) decreases differentiation rate r_H_ while also increasing the lipid production rate k_H_, and both were assumed to follow Michaelis-Menten kinetics. We also interpreted the effect of insulation by PPARγ autoregulation as a factor, I, that changed the sensitivity of HIF1α-driven negative regulation. Thus, the insulation factor controls the balance between HIF1α’s activation and inhibition effect. This factor is determined by how much PPARγ feedback is established. Under these conditions, we obtained the analytical solution for our simplified model (supplementary text). By varying the degree of insulation and HIF1α overexpression, we observed that when insulation factor, I, is low, HIF1α overexpression monotonically decreases lipid production, whereas when I is high, HIF1α overexpression increases lipid production (Figure S5B). Importantly, our simplified model also recapitulated the observation that mild HIF1α overexpression increased lipid production while severe overexpression decreased lipid production. Thus, results from both of our models strongly argued that the role change and biphasic response of HIF1α is an emergent property of the circuit’s paradoxical architecture.

### Three parameters explain a wide range of paradoxical observations

We were particularly excited to identify a biphasic lipid accumulation that is dependent on the strength of adipogenic stimulation and strength and timing of HIF1α signaling, response during differentiation, as it can explain previous reports on the effects of mild and severe hypoxia (24). We wondered if other conflicting earlier studies could also be understood in the context of the three critical factors that we identified: (1) the strength of the differentiation cue, (2) timing of hypoxic perturbation, and (3) the strength of HIF1α expression changes. For example, the strength of the differentiation differs between in vitro and in vivo systems. Studies of adipogenesis using cell culture models generally use a strong differentiation cue that generates a large amount of fat cells (10-100 %). In contrast, the apparent differentiation cue must be weaker in vivo, since previous estimates report that only about 0.5-1.65% of preadipocytes undergo differentiation at any given time (Ahrends et al., 2014; Spalding et al., 2008). Another example of differences between studies are the frequently used classical adipose promoters AP2 and adiponectin to manipulate adipogenesis in animal experiments. Because both adiponectin and AP2 are induced by PPARγ very late in adipogenesis, perturbations based on these promoters act selectively on differentiated cells, rather than by preventing the commitment step for preadipocytes which is driven earlier by autoregulation of PPARγ (Ahrends et al., 2014; Bahrami-Nejad et al., 2018)(supplementary text).

To evaluate how different published studies fit into this three-factor regulation in our model, we first generated a three-dimensional landscape relating the three factors to lipid accumulation in fat cells (Figure 5C). By including three dimensions, one can visualize how lipid accumulation is impacted by HIF1α expression and the timing of the perturbations at different strength of the differentiation stimulus (Figure 5D). We then mapped the results of thirteen different experimental studies performed by different groups (Table S2) onto this three-dimensional map. We simplify the representation into two maps for low and high differentiation stimuli, respectively. This type of analysis is relevant since the previous studies have drawn conclusions that are at odds with one another and left open the question what suitable drug therapies could minimize the HIF1α - mediated adipose tissue dysregulation (Ban et al., 2014; Gonzalez et al., 2019; Hammarstedt et al., 2018). Markedly, the mapping onto this regulatory space shows that these findings are in fact consistent with one another if the three factors of hypoxic perturbation timing, HIF1α expression and differentiation cue strength are taken into account as different conditions in the respective experiments (Figure 5E, Table S2, supplementary text).

## Discussion

To understand how cells integrate adipogenic and hypoxic cues is a daunting challenge. Numerous signaling pathways are involved, which has made it difficult to develop a consistent model. To reconcile such complexity, we adopted a combined experimental and modeling approach which enables a more complete exploration of the integrated capability of the system. Our modeling and experimental work identified a circuit that insulates two opposing roles of HIF1α on lipid accumulation in adipocytes. Before preadipocytes engage the positive feedbacks that increase PPARγ and commit the cells to irreversibly differentiate into adipocytes, a HIF1α increase blocks differentiation and lipid accumulation by suppressing the increase in PPARγ. In contrast, a HIF1α increase in differentiated adipocytes induces a metabolic switch that leads to strong upregulation of lipid accumulation which only occurs in differentiated cells with persistently high PPARγ. Thus, increases in HIF1α expression can have two opposing effects on lipid accumulation in adipose tissues that both can cause adipose dysregulation: HIF1α prevents new adipocytes to form, and thereby reduces lipid accumulation, but HIF1α also promotes accumulation of high amounts of lipids in existing adipocytes which can exceed the healthy range.

An intriguing hypothesis that emerges from our work is that the rapid increase in fat cell size (hypertrophy) in obesity may cause two normally separated, healthy roles of HIF1α to be joined in a double-negative feedback relationship that could lock adipose tissue into an unhealthy inflammatory state leading to subsequent fibrosis and insulin resistance. As shown schematically in Figure 6, the hypertrophic enlargement of fat cells suppresses oxygenation of fat, which leads to increased HIF1α activity that further promotes the development of hypertrophy. HIF1α-driven secretion of inflammatory factors is sustained as a by-product of the sustained double-negative feedback.

**Figure 6.**
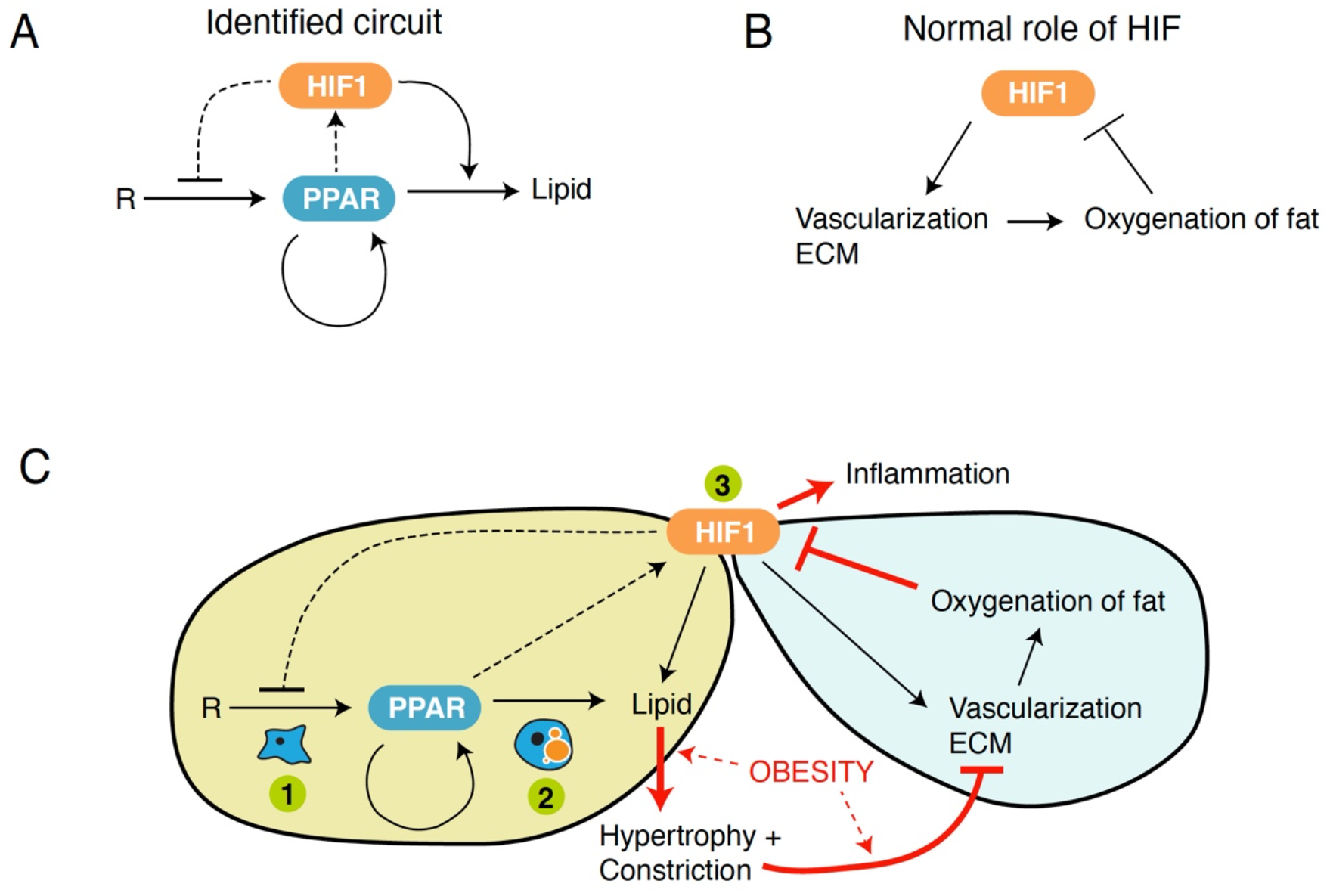
Schematic of how two normally healthy HIF1*α* circuits in adipose tissue could be locked by obesity into a sustained state of inflammation. (A) Experiments and modeling identified an adipogenic circuit in which HIF1α has two opposing roles that are isolated from each other by PPARγ autoregulation. (B) Canonical role of HIF1α to sense and maintain healthy levels of oxygen in tissues. (C) Excessive lipid accumulation and expansion of adipocyte size (hypertrophy) in obesity increases oxygen consumption and suppresses vascularization in adipose tissue. The reduction in oxygenation of the fat causes HIF1α to remain high, which in turn drives more suppression of adipogenesis (green circle #1) and more lipid accumulation into mature fat cells (green circle #2). Both of these latter effects cause more adipocyte hypertrophy and more constriction of vasculature, leading HIF1α to be locked into a continually high state. A sustained HIF1α state is harmful since, whereas short-term HIF1α expression has beneficial effects to restore oxygen homeostasis, HIF1α controls the expression of many inflammatory and ECM remodeling genes which in the long-term causes fibrosis and insulin resistance. Because the PPARγ autoregulation isolates an opposing signaling role of HIF1α’s before and after differentiation commitment, therapies to revert the harmful sustained HIF1α state that only promote differentiation of preadipocytes (green circle #1), suppress lipid accumulation of adipocytes (green circle #2), or inhibit HIF1α (green circle #3) are not predicted to be effective on their own. In contrast, a simultaneous targeting at all three green points has the potential to reverse the HIF1α-driven disease state (all three green circles).

Since our model shows that the roles of HIF1α in pre-adipocytes and adipocytes can be isolated, synergistically manipulating the balance between the positive and negative roles would be a novel strategy for disrupting the unhealthy double-negative feedback brought on by obesity. For example, our model suggests that a three-point strategy combining a PPARγ agonist, like rosiglitazone, to increase adipogenesis (#1, Figure 6C), reducing lipid uptake into adipocytes, possibly with a CD36 inhibitor (#2, Figure 6C), and reducing HIF1α activity with a HIF1α inhibitor (#3, Figure 6C) could improve metabolic outcomes by effectively promoting adipogenesis and reducing adipocyte hypertrophy in a synergistic manner while still allowing for lower, healthy levels of HIF1α activity. Such synergistic multi-point therapies may be effective as suggested by observed clinical improvements from a previous study combining rosiglitazone and metformin in treating type 2 diabetics (Zinman et al., 2010).

In summary, we describe an integrated experimental and modeling approach to investigate HIF1α signaling in the context of adipogenesis and lipid synthesis. By accounting for a time-dependent dual role of HIF1α, our approach yielded several surprising and experimentally verified predictions. Most notably, we found a positive-feedback mediated insulation separates two opposing functions of HIF1α in controlling adipose mass. By incorporating the dual role of HIF1α into a computational model, we could explain a variety of contradictory results in the literature based on differences in experimentally controllable parameters. These results expand our knowledge of basic adipose tissue biology and suggests potential future drug combination treatment strategies of disorders associated with adipose dysfunction.

## Materials and Methods

### Cell culture and differentiation

Cells were maintained in MEM-ɑ media (Invitrogen, #12561) supplemented with 20% Fetal Bovine Serum (FBS), 100 units/mL Penicillin, 100 ug/mL Streptomycin, and 292 ug/mL L-glutamate (Invitrogen, #10378-016) at 37 °C, 5% CO_2_. To induce differentiation, either (1) rosiglitazone or (2) DMI protocol was used. OP9 cells were plated at 10,000 cells/well on a 96-well glass-bottom plate. The following day, the media was replaced with MEM-ɑ media supplemented 10% FBS plus (1) rosiglitazone or (2) 0.517 mM IBMX (Sigma Aldrich, USA), 62 nM dexamethasone (Sigma Aldrich, USA) and 172 nM insulin (Sigma Aldrich, USA). After two days, the media was replaced with MEM-ɑ media supplemented with 10% FBS and 172 nM insulin every two days for two or four more days. For hypoxic experiments, cells were cultured in another incubator (BINDER CO2 incubators CB-150; Binder Scientific) at 37 °C, 1% O_2_, 5% CO_2_. Doxycycline (Millipore Sigma, D9891) was used to induce dnCEBP and tre-stblHIF.

### Generation of stable cell lines

For the genetic perturbation, we lentivirally delivered the following constructs to OP9 cells.

Cells were then selected by antibiotic selection or sorting using a fluorescent marker.

- dnCEBP: pLenti-TRE-A-C/EBP-PGK-rtTAA-2A-puro
- stblHIF (for OP9^HIF^): pLenti-PGK-stblHIF-PGK-puro
- tre-stblHIF (for OP9^treHIF^): pLenti-TRE-stblHIF-PGK-rtTAA-2A-puro tre-stblHIF codes HIF1α with triple mutations (P402A, P577G and N813A) as previously described (*30, 31*). These constructs will be available from Addgene.

### Immunofluorescence

OP9 cells were fixed with 4% paraformaldehyde in PBS for 15 min. Then the cells were washed three times with PBS. Permeabilization was carried out with 0.1% Triton X-100 in PBS for 15 min, followed by blocking with 5% bovine serum albumin (Sigma #7906) or 10% donkey serum (Jackson Laboratory). The cells were stained with anti-PPARγ (1:1000 Santa Cruz Biotech #sc-7273) and/or anti-HIF1αlpha (1:800 Santa Cruz Biotech #sc-7273 or 1:800 Cell Signaling Technology #36169S). For secondary antibodies, FITC-conjugated anti-mouse antibody (Jackson Laboratory #115-095-166), Rhodamine-conjugated anti-rabbit antibody (Jackson Laboratory, #111-295-144), or FITC-conjugated anti-rabbit antibody (Jackson Laboratory, #111-095-144) were used. After washing secondary antibodies, cells were further stained with Hoechst 33342 (50 ng/mL) and LipidTox Deep Red Neutral Lipid Stain (1:20,000 Thermo Fisher Scientific). As the lipid droplets slowly diffused over time, LipidTox stain and imaging were conducted within a day after fixation.

### Time-lapse microscopy

A day prior to imaging, OP9 cells expressing PPARγ-mCitrine were plated at a density 10,000 cells/well on a fibronectin-coated (10ug/ml) (Sigma Aldrich, F0895) 96-well glass imaging plate (Fisher Scientific, 164588). To initiate the differentiation, the growth media was replaced with 200uL of imaging media (FluoroBrite DMEM (Thermo Fisher Scientific, A1896701) supplemented with 292 ug/mL L-glutamate and 10% FBS) containing 250 *µ*M rosiglitazone. An AeraSeal film (Sigma Aldrich, A9224) was applied to the plate to prevent evaporation. Imaging was performed using Nikon Eclipse Ti fluorescence microscope with Andor Neo 5.5 sCMOS controlled by Micromanager, equipped with temperature (37°C) and environmental control (5% CO_2_). Images were acquired at 20-minute intervals using 10x/0.30 numerical aperture objective and 1.5x tube lens at 3×3 binning. 48 h after the differentiation, the media was replaced with the fresh imaging media containing either DMSO or 50 *µ*M FG-4592.

### Image analysis

Image processing and analysis were performed using CellTK, a custom software developed in Python (https://github.com/CovertLab/CellTK) (Kudo et al., 2018). For analysis of stained images, after illumination correction and background subtraction, fluorescent images of nuclear markers were segmented by thresholding of the Laplacian of Gaussian of the image followed by an adaptation of the morphological snakes algorithm to round out the nuclei. Then cytoplasmic masks were created by the dilation of nuclei a set number of pixels followed by the exclusion of background pixels to generate a ring-like segmentation around nuclei. Median fluorescence intensities of the pixels corresponding to the nuclear region (for PPARγ and/or HIF1α) and cytoplasmic region (for lipid staining) in each individual cell were measured for each fluorescent channel. Mean intensity was then calculated from > 2,000 cells per technical replicate. We note that measurements of cytoplasmic lipid intensity at the single-cell level are noisier than nuclear PPARγ and HIF1α as our cytoplasmic segmentation only accounts for lipid droplets around nuclei. For the live-cell image analysis, the segmented nuclei were further tracked from frame-to-frame by primarily using a linear assignment problem (LAP) based approach. The remaining cells that were not captured by this method were subject to watershed separation followed by nearest neighbor tracking.

### Knockdown experiment

OP9 cells were transfected with the respective siRNA or control siRNA 24 hours before the start of an experiment by reverse transfection using Lipofectamine RNAiMax (Invitrogen #13778150), following the manufacturer’s protocol. The following siRNA was used as a pool: PPARγ (Qiagen SI01385391, SI01385398, SI01385405), CEBPA (Qiagen SI00948311, SI00948318, SI00948325), CEBPB (Qiagen SI00948339, SI00948360, SI04423083), FABP (horizon L-042923-01-0005) and HIF1α (Qiagen SI00193032, SI00193025, SI00193018, SI00193011). siHIFv1 and siHIFv2 correspond to two individual siRNA, SI00193032 and SI00193025, respectively. For negative control siRNA, AllStars Negative Control siRNA (Qiagen #1027280) or ON-TARGETplus Non-targeting siRNAs (horizon, D-001810-0X) were used.

### Measurement of glycolytic flux

100 *µ*g/ml poly-d-lysine (Sigma, A-003-M) was added to each well in the Seahorse XF96 V3 PS Cell Culture Microplates and incubated for two hours, followed by three times wash with H_2_O. After one hour, OP9 cells were transfected with siRNA for HIF1α or control siRNA and plated at 6,000 cells per well in the poly-d-lysine coated plate. The next day, OP9 cells were differentiated according to the rosiglitazone differentiation protocol but using DMEM with 1 mM sodium pyruvate, 2 mM glutamine, 5.5 mM glucose, 10% FBS, and 3.7 g/l sodium bicarbonate. On day 4 after inducing differentiation, glycolytic flux measurements were performed using a Seahorse XF Glycolytic Rate assay protocol on a XF96 extracellular flux analyzer (Seahorse Biosciences). The assays were conducted in the media without sodium bicarbonate and FBS. Inhibitors were injected to final concentrations of 0.5 *µ*M antimycin A (Sigma Aldrich, USA), 0.5 *µ*M rotenone (Sigma Aldrich, USA), and 100 mM 2-Deoxy-D-glucose (2DG) (Sigma Aldrich, USA). After the Seahorse assays, the cells were fixed and stained with Hoechst as described above and imaged. To account for the changes in cell number during differentiation, the measured extracellular acidification rate (ECAR) and oxygen consumption rate (OCR) in each well were normalized by the number of cells, obtained by counting the nuclei in the Hoechst stained images. The buffer factor in the Glycolytic Rate assay was estimated by fitting the glycolytic proton efflux rate after 2DG treatment to 0.

### Statistical Analysis

Statistical analysis was carried out in Python and a description is available in the main text, method details and associated figure legends.

## Acknowledgments

We thank Tobias Meyer, Zhi-bo Zhang, and Agnes Agas (Weill Cornell) for discussions and for providing feedback on the manuscript. Sorting was performed on an instrument in the Shared FACS Facility obtained using NIH S10 Shared Instrument Grant (S10RR025518-01).

## Author contributions

Conceptualization: MLZ, TK

Methodology: EL, KK, MLZ, TK

Investigation: MLZ, TK

Visualization: TK

Software: TK

Funding acquisition: MNT, MWC

Project administration: MLZ, TK

Supervision: MNT, MWC

Writing – original draft: MLZ, TK, MNT, MWC

Writing – review & editing: EL, MLZ, TK, MNT, MWC

## Data and materials availability

Simulation source code will be available on GitHub (https://github.com/braysia/hifppar). Further resources and reagents will be available upon request.

## Supplemental Information

Figure S1. HIF1α regulates glycolytic flux in differentiating OP9 cells. Related to Figure 1.

Figure S2. Adipogenic induction of HIF1α is stimulus-independent. Related to Figure 2.

Figure S3. PPARγ-HIF1α model simulation results were consistent with multiple experimental observations. Related to Figure 3.

Figure S4. Differentiation cue-dependent lipid accumulation response under conditions of HIF1α inhibition. Related to Figure 4.

Figure S5. Biphasic response is an emergent property of a paradoxical architecture.

Table S1. Description of model parameters.

Table S2. A list of in vivo and in vitro studies.

