## Supplemental Material for "A dynamic HIF1α- PPARγ circuit controls a paradoxical adipocyte regulatory landscape"

**This PDF file includes:**

Supplementary Text

Figures S1 to S5

Tables S1 to S2

**Supplementary Text**

<u>Model architecture</u>

We elaborate on our previous model with PPARγ driven positive feedback (Ahrends et al., 2014) and further include HIF1α and lipid regulation. In this study we found both negative feedback and positive feedforward of HIF1α play a substantial role in overall lipid accumulation. Our aim is to study how this architecture with paradoxical regulation contributes to the cellular outcomes when perturbed under various scenarios. We note that our focus was to coarsely model the relationship between PPARγ and HIF1α, and the detailed molecular mechanisms remain unknown. However, we provide further clarification for our model.

$$\frac{d[PPAR]}{dt} = \varepsilon \cdot k_1 \cdot \left( [R] \cdot k_r \cdot \frac{[CEBP]^3}{K_{inh}^3 + [CEBP]^3} \right) - d_p \cdot [PPAR] \qquad (1)$$

$$\frac{d[CEBP]}{dt} = k_2 \cdot [PPAR] - d_c \cdot [CEBP] \qquad (2)$$

$$\frac{d[HIF]}{dt} = k_3 \cdot \frac{[PPAR]}{K_H + [PPAR]} - d_H \cdot [HIF] \qquad (3)$$

$$\frac{d[LIPID]}{dt} = k_4 \cdot [HIF] \cdot \frac{[PPAR]^3}{K_H^3 + [PPAR]^3} - d_L \cdot [LIPID] \qquad (4)$$

$$K_{inh} = K_C \cdot \left( 1 + \frac{[HIF]}{K_i} \right) \qquad (5)$$

Equation (1 & 2): PPARγ synthesis in the model is controlled by R and the positive feedback from CEBP. The R parameter is intended to be a ligand activating PPARγ, which mimics the effects of rosiglitazone. Although the physiological ligand for PPARγ is unknown, the precise pathways responsible for the initial activation of PPARγ are not important for our model. PPARγ has a number of positive feedback partners, but here we compiled them into one strong nonlinear Hill equation represented by CEBP. One mode of negative regulation of PPARγ by HIF1α was previously characterized to be mediated through a transcriptional repressor BHLHE40 (Yun et al., 2002). BHLHE40 binds to the upstream region of PPARγ promoter and represses transcription. Thus we modeled this regulation as a competitive inhibition term that limits the transcription from CEBP; however, we note that there may be other factors involved in this process, including the recently discovered PPARγ phosphorylation via the HIF1α-ERK pathway (*3*) or post-translational modification of PPARγ coactivator 1a (Krishnan et al., 2012). For each simulation, a noise factor $\boldsymbol{\varepsilon}$ was sampled from a log-normal distribution with sigma set to 0.3 to represent cellular variation.

Equation (3): We also assume the presence of endogenous HIF1α, according to our experimental model. HIF1α level during adipogenesis was controlled by PPARγ (Figure 2C and Figure S2A). Our data and data in the literature suggests that differentiation ,which is caused by a strong increase in the transcriptional output of PPARγ, in turn promotes HIF1α expression which is a main regulator of "active" HIF1α production. (Figure 1C and 1G). Biologically, HIF1α is primarily regulated through degradation; here, in order to minimize the model complexity, we simply assumed the rate of "active" HIF1α production being proportional to the transcriptional output of PPARγ, which is expressed as Michaelis-Menten kinetics.

Equation (4): The production term of lipid is modeled as a product of functions of HIF1α and PPARγ. The lipid production is mostly linear to HIF1α as we showed in Figure 1D. The dependency on PPARγ is modeled as a Hill equation, to incorporate a time delay in the system. Biologically, this integrates many functions of PPARγ such as HIF1α independent metabolic regulation including GLUT4 induction. For the implementation, see https://github.com/braysia/hifppar. Unless noted, we visualized the endpoint simulation values (at day 6 after differentiation was induced).

<u>Simulation of perturbation</u>

Overexpression in the model refers to the overexpression using genetic construct, activation through chemical activators, or hypoxia. HIF1α expression caused by hypoxia was additive to the expression induced by PPARγ (Figure S3G). Given that the experiment was performed at a saturating amount of rosiglitazone, we consider HIF1α induction by hypoxia to be a part of an independent pathway. To model overexpression, we thus give an additional HIF1α in a step-function manner. We add a term $E_{ov}(t)$ for overexpression to modify equation (4) and (5):

$$\frac{d[LIPID]}{dt} = k_4 \cdot \left( [HIF] + E_{ov}(t) \right) \cdot \frac{[PPAR]^3}{K_H^3 + [PPAR]^3} - d_L \cdot [LIPID]$$

$$K_{inh} = K_C \cdot \left( 1 + \frac{[HIF] + E_{ov}(t)}{K_i} \right)$$

where

$$E_{ov}(t) = \begin{cases} HIF_{over} \cdot HIF_{SS} & \text{if } t > \tau \\ 0 & \text{if } t \leq \tau \end{cases}$$

$\tau$ controls the timing of overexpression and $HIF_{over}$ denotes the strength of overexpression. When $HIF_{over} = 1$, 0.025 µM HIF1α (the steady-state when the system is stimulated with R=1.0) is introduced as an overexpression.

Similarly, to model HIF1α knockdown or inhibition, we introduce the term $E_{inh}(t)$ to equation (3):

$$\frac{d[HIF]}{dt} = E_{inh}(t) \cdot \left( k_3 \cdot \frac{[PPAR]}{K_H + [PPAR]} \right) - d_H \cdot [HIF]$$

where

$$E_{inh}(t) = \begin{cases} 1 + HIF_{kd} & \text{if } t > \tau \\ 0 & \text{if } t \leq \tau \end{cases}$$

$\tau$ controls the timing of inhibition, $HIF_{kd}$denotes the strength of inhibition. $HIF_{kd}$takes a value ranging from 0 to -1, where -1 indicates a complete inhibition or knockout.

To compromise the PPARγ feedback, we chose to use a dominant-negative form of CEBP. To represent its effect in the model, we simply limit the contribution of CEBP to PPARγ by introducing $C_i$.

$$\frac{d[PPAR]}{dt} = \varepsilon \cdot k_1 \cdot \left( [R] \cdot k_r \cdot \frac{C_i \cdot [CEBP]^3}{K_{inh}^3 + C_i \cdot [CEBP]^3} \right) - d_p \cdot [PPAR]$$

We chose $C_i$ to be 0.33 (equivalent to having one-third of CEBP) (Musunuru et al., 2010; Olive et al., 1996; Zhang et al., 2004).

We also wanted to test if the paradoxical dual action of HIF1α is required for the role change of HIF1α overexpression over time in lipid accumulation. Thus we modified the model structure to abrogate (1) inhibition of PPARγ synthesis, and (2) induction of lipid synthesis, by HIF1α.

(1) The "negative only" model uses the following for equation (5):

$$K_{inh} = K_C$$

(2) The "positive only" model uses the following for equation (4):

$$\frac{d[LIPID]}{dt} = k_4 \cdot \frac{[PPAR]^3}{K_H^3 + [PPAR]^3} - d_L \cdot [LIPID]$$

For each model, we performed the randomized parameter search described in Table S1 by sampling random values for free parameters 30,000 times. We then recorded the fraction of the parameter set that could recapitulate the timing dependent effect to any extent by setting the following criteria: (1) early overexpression (from 0 h) of HIF1α led to >1% reduction of lipid level; (2) later overexpression (from 48 h) of HIF1α led to >1% increase of lipid level.

<u>Randomized parameter search to obtain free parameters</u>

The free parameters, $k_2$, $K_C$, $K_H$, $K_i$, were chosen to reproduce the timing-dependent effect through randomized search. For each simulation, we randomly pick a parameter set from a grid ranging from $10^{-5}$ to $10^1$ in log-scale. We calculate the sum of lipids from 200 cells using R=0.25 and$HIF_{over}$=1 with varying $\tau$. $k_1$and $k_4$were computed according to each parameter set. This process was repeated 30,000 times. The parameter candidates were first selected based on the lipid level at 96 h to represent the paradoxical timing-dependency:

(1) The lipid level decreases to 50±25% when overexpression at $HIF_{over}$=1 was induced from 0 h at R=0.25.
(2) The lipid level increases to 200±50% when overexpression at $HIF_{over}$=1 was induced from 48 h at R=0.25.

Using the selected candidates, we repeated the simulation using R=1.0 and R=0.001 (assuming the saturating and basal condition, respectively). We then chose the final set of parameters such that they minimize the following objective, in order to fulfill some of the steady-state assumptions used above:

$$\text{minimize}\left|\left(\frac{[HIF_{R=1}]}{HIF_{SS}}-1\right)^2+\left(\frac{[PPAR_{R=1}]}{PPAR_{SS}}-1\right)^2+\left([LIPID_{R=0.001}]\right)^2\right|$$

Description of population dynamics model

We assume cells can take one of the two states of PPARγ, either $PPAR_{low}$ (x) or $PPAR_{high}$ (y). The fraction of each population is described by the following ODE, where differentiation from $PPAR_{low}$ to $PPAR_{high}$ is determined by a differentiation rate $r_H$. We simply assume the total lipid level (z) is a product of synthesis rate ($k_H$) and the fraction of PPARγ high cells (y).

$$\frac{\mathrm{d}x}{\mathrm{d}t}=y-r_H\cdot x$$

$$\frac{\mathrm{d}y}{\mathrm{d}t}=r_H\cdot x-y$$

$$z=k_H\cdot y$$

To incorporate HIF1α's dual action to this model, we assume HIF1α (H) negatively affects the differentiation rate $r_H$ and positively impacts the synthesis rate $k_H$. We modeled these contributions as Michaelis-Menten equations:

$$r_H=1-\frac{H}{K_d+H}$$

$$k_H=1+\frac{H}{K_l+H}$$

Without HIF1α overexpression, both $r_H$ and $k_H$ are set to 1. Solving these equations for y,

$$y(t)=\frac{r_H}{r_H+1}-r_H\cdot\frac{e^{-(r_H+1)t}}{r_H+1}$$

Then the steady state adipocyte fractions can be expressed as

$$y_{ss} = \frac{r_H}{r_H + 1} = \frac{K_d}{H + 2K_d}$$

The steady state lipid level is then

$$z_{ss} = k_H \cdot y_{ss} = \frac{K_d\left(K_l + 2H\right)}{\left(K_l + H\right)\left(2K_d + H\right)}$$

The relative changes in lipid by HIF1α overexpression is expressed as

$$\frac{z_{ss}}{z_{ss}^{H=0}} = \frac{2K_d\left(K_l + 2H\right)}{\left(K_l + H\right)\left(2K_d + H\right)}$$

Now we simply assume insulation (*I*) as a factor to control the sensitivity of negative regulation ($K_d$) relative to the sensitivity of positive regulation ($K_l$). To find out the range of HIF1α overexpression which works positively to the lipid,

$$K_d = I \cdot K_l$$

$$\frac{z_{ss}}{z_{ss}^{H=0}} > 1$$

$$H^{+} < 2K_d - K_l = K_l(2I - 1)$$

Thus, HIF1α overexpression monotonically decreases the lipid level when I < 0.5.

Mapping previous studies to the regulatory landscape

Here we describe how we mapped the previous studies to the landscape (Figure 5; Table S2). In the studies conducted with mice, the apparent differentiation cue, R, is considered low as the turnover rate is slow *in vivo*. The differentiation rate was previously estimated to be 0.5 - 1.65 %, of which the corresponding R-value in our model becomes roughly 0.004-0.005 in our simulation. In contrast, we consider all the experiments conducted in cell culture to have a high differentiation cue. In most studies in cell culture, the differentiation rate is an order of magnitude higher than that *in vivo* (10-100%).

Generally, adipose-specific perturbation in mice uses Cre recombinase under the control of a promoter driven by PPARγ. Thus we consider these systems to be in the late phase, except for the study from Zhang et al (Zhang et al., 2010). Zhang et al (Zhang et al., 2010) and Sun et al (Sun et al., 2013) used the same dominant-negative construct to restrict HIF1α activity, yet the latter is considered to have a tighter control in the timing of expression as it uses the doxycycline-inducible promoter. The former study uses a mouse model reported to be leaky with about 40% of undifferentiated cells in the stromal vascular fraction already expressing ap2 promoter (Shan et al., 2013). Thus, we classified the study from Zhang et al to be in the early phase with respect to Sun et al. To further improve our classification, the actual timing of perturbation needs to be further characterized (Jeffery et al., 2014; Shan et al., 2013). We also classified the in vivo administration of small molecule inhibitors (Sun et al., 2013) or antisense oligo (Shin et al., 2012) as occurring in the late phase, since a large portion of adipogenesis appears to occur early in the development in mice (Wang et al., 2013). Although further studies are required to determine how the slow turnover rate in humans (Spalding et al., 2008) would impact a longer-term effect, the overexpression of HIF1α is likely to reduce the fraction of mature adipocytes. This interpretation is supported by observations of the increased number of small, immature adipocytes under conditions of insulin resistance, a disorder closely associated with dysregulated HIF1α signaling (McLaughlin et al., 2007).

Most in vivo experiments were highly biased against the “early” perturbation due to the limited molecular tools. We classified the study by Rahtu-Korpela et al (Rahtu-Korpela et al., 2014) as early, as it is not an adipose-specific perturbation. Due to the other functions of HIF1α during development, however, the mechanism leading to the weight reduction in such a system is possibly attributed to a more systemic effect (Thomas et al., 2016). The recent identification of the pre-adipocyte promoter could be useful to test the hypothesis we generated in this study (Jeffery et al., 2014). On the other hand, we classified the current *in vitro* model using 3T3-L1 cells biasing toward the “early” perturbation; the use of late perturbation in cell culture may produce more consistent results with in vivo Cre recombinase model.

One limitation to note is that our sampling of the landscape is partially arbitrary. Slight differences in R, timing or degree of perturbation may lead to slightly different conclusions with respect to our mapping of previous studies. For example, we chose R=0.01 as a low R in Figure 5D, but the apparent R-value may be even lower given the reported low *in vivo* differentiation rate. However, our choices in parameter values are intended to serve an exemplary purpose and demonstrate the sensitivity of the system to small changes. To further highlight this point we note that the same ap2-Cre system used to generate the adipose-specific PhD2 knockout by Matsuura et al (Matsuura et al., 2013) and Michaelidou et al (Michailidou et al., 2015) resulted in opposite outcomes. Based on our model, this apparent contradiction may be attributed to differences in important factors such as the degree of knockout in the adipocyte population and the timing of knockout based on the activation of the Ap2 promoter. These systematic variations are likely to exist since the ap2-Cre system has been reported to have variable efficacy when used for the targeting of adipocyte cell populations (Jeffery et al., 2014; Shan et al., 2013).

### Supplementary Figures

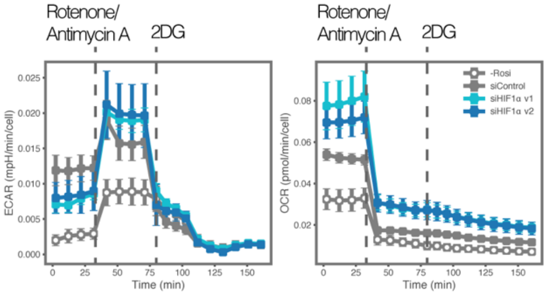


**Figure S1. HIF1α regulates glycolytic flux in differentiating OP9 cells.** HIF1α knockdown significantly reduced the rate of glycolytic flux. The flux was measured using the Seahorse Glycolytic Rate Assay. Extracellular acidification rate (ECAR, left panel) and oxygen consumption rate (OCR, right panel) upon rotenone plus antimycin treatment A followed by 2DG treatment were measured to calculate the glycolytic rate. ECAR and OCR values were normalized by the number of cells obtained from the Hoechst-stained images after the assay. Cells were differentiated for four days (1 μM rosiglitazone) with or without HIF1α siRNA transfection. Line and bar plots are presented as means ± SD from four replicates.

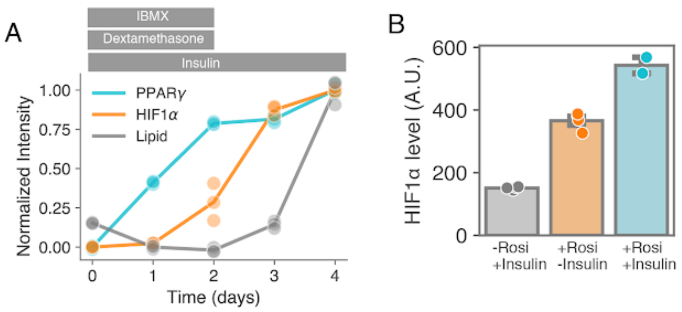


**Figure S2. Adipogenic induction of HIF1α is stimulus-independent.**
(**A**) HIF1α was induced by DMI stimulation in a similar manner as was observed using rosiglitazone induction. OP9 cells were differentiated using a standard DMI cocktail. After 48 hours the cocktail was removed and replaced with insulin alone.
(**B**) HIF1α induction is enhanced but does not require the addition of insulin into the media. Media containing rosiglitazone was removed after 48 hours of differentiation induction and replaced with media with or without insulin.

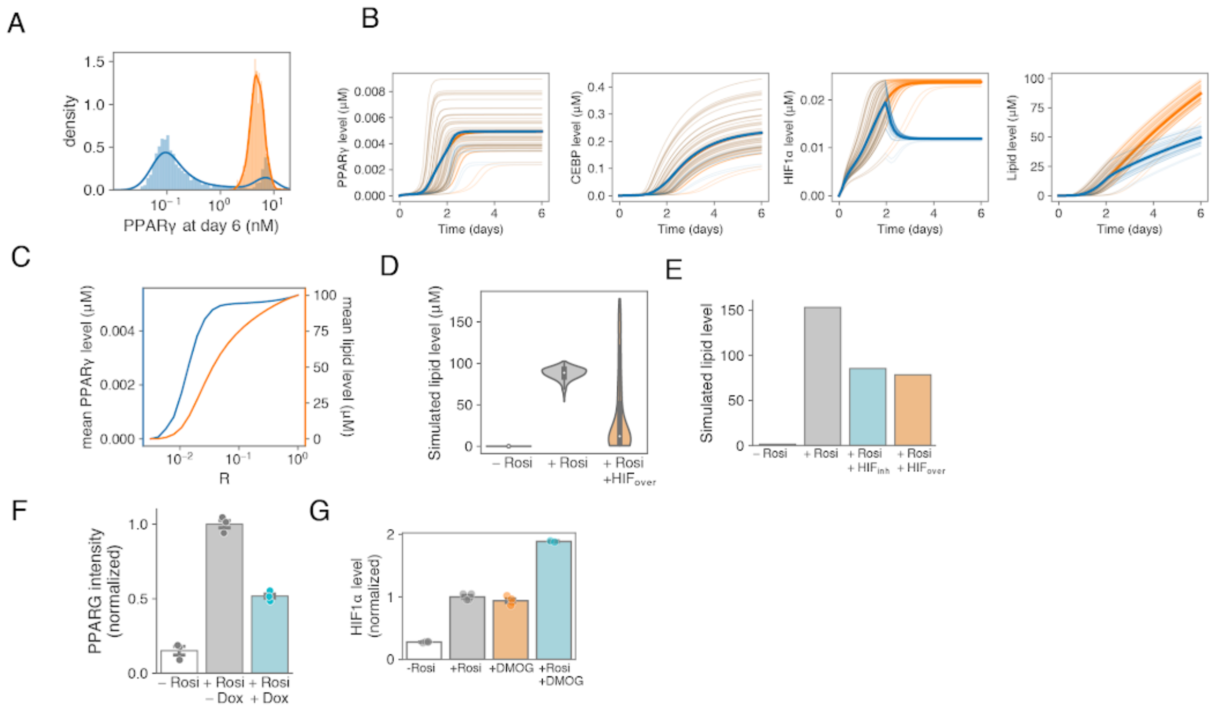


**Figure S3. PPARγ-HIF1α model simulation results were consistent with multiple experimental observations.**

(**A**) Bistable behavior of PPARγ. PPARγ levels were simulated at R=0.25 with (blue, $HIF_{over}$=1) or without (orange) HIF1α overexpression. The normalized histogram was plotted using 1000 cells for each condition.

(**B**) Representative time-traces. The model simulates the time-course of PPARγ, CEBP, HIF1α, and lipid concentration over 6 days. Cells were simulated at R=0.25 with (blue, $HIF_{kd}$=0.5) or without (orange) HIF1α knockdown. Representative time-traces of 50 cells and their mean (solid line) were shown.

(**C**) A dose-response curve of PPARγ and lipid concentration.

(**D**) The simulation of HIF1α overexpression reproduced the long-tail distribution of lipid concentration. The lipid level was simulated at R=0.001 (-Rosi) or R=0.25 (+Rosi) with (orange) or without HIF1α overexpression at $HIF_{over}$=1. The simulated single-cell distribution of lipid level at day 6 was plotted with a box plot showing the mean and quantiles.

(**E**) The simulation reproduced the reduced lipid level by both HIF1α knockdown and overexpression. The lipid level was simulated at R=0.001 (-Rosi) or R=0.25 (+Rosi), combined with HIF1α knockdown (cyan, $HIF_{kd}$=0.5) or overexpression (orange, $HIF_{over}$=1).

(**F**) Doxycycline treatment reduced the PPARγ level at day 4 in dnCEBP OP9 cells. Cells were differentiated (1 μM rosiglitazone for +Rosi) and treated with 0.4 μg/ml doxycycline (for +Dox). Each dot represents a biological replicate and the bar plot is presented as means ± SD.

(**G**) HIF1α induction by the hypoxic mimetic DMOG was additive to rosiglitazone-induced HIF1α. A day prior to fixation, DMOG (1 mM) was added to a typical four-day rosiglitazone differentiation (1 μM). HIF1α levels were then assessed by immunofluorescence after day 4. Each dot represents a biological replicate and the bar plot is presented as means ± SD.

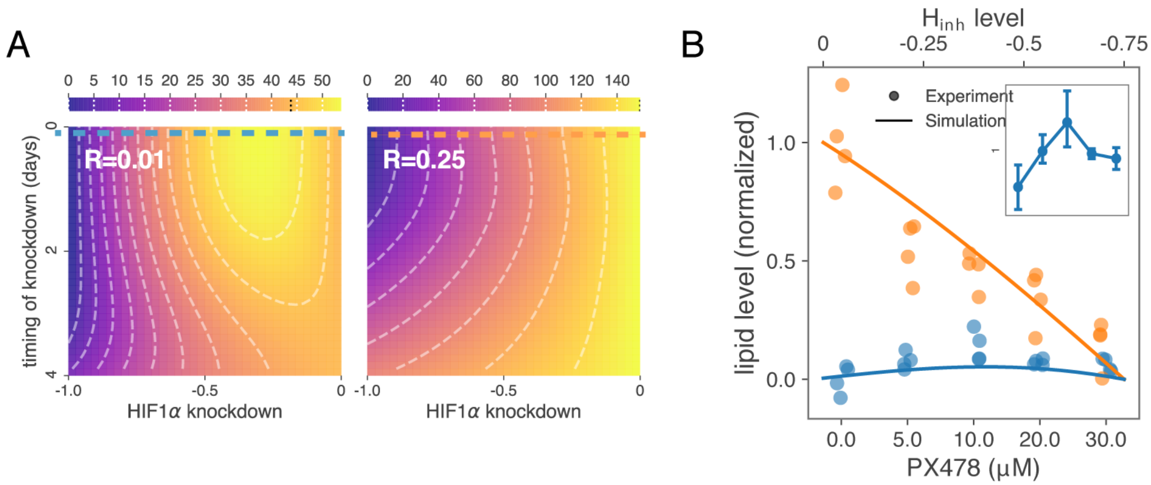


**Figure S4. Differentiation cue-dependent lipid accumulation response under conditions of HIF1α inhibition.**

(**A**) Model simulations predict that a biphasic response also occurs in the regime of HIF1α inhibition. Each colored plot shows the simulated degree of lipid synthesis for various HIF1α knockdown levels and timing of HIF1α inhibition. The left plot shows the simulation results for a relatively weak differentiation cue while the right plot shows the results for a comparatively stronger differentiation cue. Contour lines are drawn as white dashed lines. The color range in the plots represents lipid synthesis values and is scaled differently for the purpose of clarity. Plots represent results from the simulation of 5000 cells for each condition.

(**B**) Two curves, the blue and orange dashed lines in (A), show qualitatively different responses to HIF1α inhibition depending on the strength of the differentiation cue. This result was validated by experiments using the small molecule PX478 to inhibit HIF1α activity in OP9 cells throughout the entire differentiation time course. Lipid levels in differentiated OP9 cells for each condition were assessed using LipoTox DeepRed staining on the fourth and final day of the differentiation protocol. Measured fluorescence values (dots) and model simulation values (solid line) were normalized to their min and max value. Orange represents high differentiation cue condition (250 nM rosiglitazone) and blue represents a low differentiation cue condition (1 nM rosiglitazone). Inset shows the close-up experimental results for the low differentiation cue showing a biphasic response, presented as means ± SD.

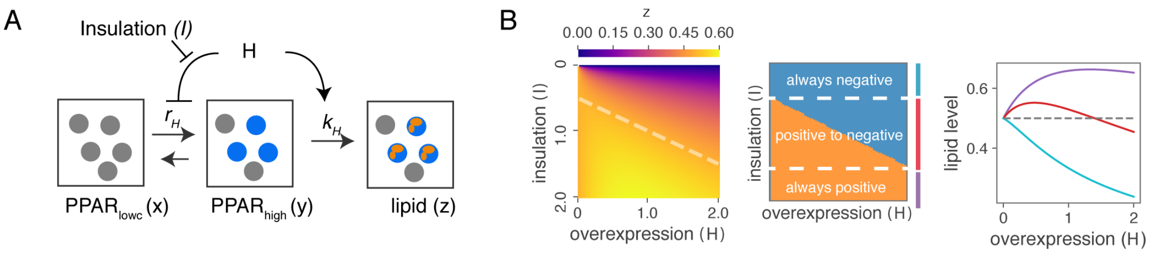


**Figure S5. Biphasic response is an emergent property of a paradoxical architecture.**
(**A**) Schematics of population dynamics model.
(**B**) (left heatmap) The lipid level landscape varying overexpression level and insulation level. Color gradients indicate the lipid level obtained from the analytical solution. $K_I$ is set to 1. The dashed line indicates the boundary between the negative and positive outcomes relative to the lipid level without overexpression, corresponding to H=2I-1. (middle panel) The insulation changes the role of overexpression. The colored region indicates a relative increase (orange) or decrease (blue) of lipid level over the condition without HIF1α overexpression. The regions separated by dashed lines show three cases of how overexpression affects the lipid level. (right panel) The population dynamics model reproduced the role change of HIF1α. Three representative curves were plotted for different insulation levels (I=0.4, 1.2 and 3.6 for cyan, red, and purple). The dash line indicates the lipid level without HIF1α overexpression.

**Table S1. Description of model parameters.**

| Parameter | Value | Description |
|---|---|---|
| $PPAR_{SS}$ | 0.005 | We assume the steady-state PPARγ level ($PPAR_{SS}$) at a saturating amount of differentiation cues (in our simulation, we set R to be 1.0 μM) to be 0.005 μM (roughly 10,000 copies per cell) (Simicevic e al., 2013). |
| $HIF_{SS}$ | 0.025 | HIF1α could go up to roughly 25 pg/mg of protein (Heikal et al., 2018). Assuming the protein conter is 300 pg/cell, this is equivalent to 0.025 μM (7.5 fg/cell). Thus, we assumed that the steady-state HIF1α ($HIF_{SS}$) is 0.025 μM at a saturating amount of differentiation cue. |
| $k_r$ | 15 | We set this value to 15 from the previous study (Ahrends et al., 2014). |
| $d_p$ | 0.5 | Assuming the half-life of PPARγ to be roughly 1.4 h (*36*), $d_P$is determined by $ln(2)/1.4$. |
| $d_C$ | 0.028 | There exist several feedback partners with different half-lives ranging from a few hours to > 30 h (Ahrends et al., 2014; Bahrami-Nejad et al., 2018). Here we represent one feedback component's hal life as 24 h. $d_C$is determined by $ln(2)/24$. |
| $d_H$ | 0.17 | Assuming the half-life of active HIF1α to be roughly 4 h (Kong et al., 2007), $d_H$is determined by $ln(2)/4$. |
| $d_L$ | 0.005 | Lipid is slowly accumulating over (possibly more than) 6 days, so we chose $d_L$to be an order of magnitude slower than the other products. |
| $k_3$ | 0.0425 | For simplicity, we assumed the steady-state level of PPARγ being greater than $K_H$ ($PPAR_{SS} >> K_H$) then $k_3$is determined by $HIF_{SS} * d_H$. |
| $k_4$ | 44 | $k_4$ is adjusted to produce 100 μM of lipid at day 6 at a saturating amount of rosiglitazone (R=1) (Campos et al., 2018). We note that this value was chosen at last since it has a minimal impact on ou parameter selection scheme that uses the ratiometric value. |
| $k_1$ | 0.000155 | For simplicity, we assumed the feedback from CEBP saturates ($CEBP_{SS}$ >> $K_C$) at a saturating amou of rosiglitazone. $k_1$ is estimated to be $d_p PPAR_{SS}(K_P + HIF_{SS})/((1 + k_r) * K_P)$. |

| | | |
|---|---|---|
| $k_2$ | 1.39 | See “Randomized parameter search to obtain free parameters”. |
| $K_C$ | 0.00209 | |
| $K_H$ | 0.000237 | |
| $K_i$ | 0.00306 | |

**Table S2. A list of in vivo and in vitro studies.**

| | Lipids | System | Perturbation | Timing | Number assigned in a figure |
|---|---|---|---|---|---|
| **HIF1 downregulation** | | | | | |
| Jiang et al (Jiang et al., 2011) | Body weight and AT mass ⇓ | aP2-Cre in C57BL6 mice | HIF1α knockout | late | 1 |
| Krishnan et al (Krishnan e al., 2012) | Body weight and AT mass⇓ | aP2-Cre-ERT2 in C57BL6 mice | HIF1α knockout | late | 1 |
| Sun et al (Sun et al., 2013) | Body weight and AT mass ⇓ | C57BL6 mice administration | PX-478 treatment | late | 2 |
| Shin et al (Shin et al., 201[illegible] | Body weight and AT mass ⇓ | C57BL6J mice administration | Antisense oligo against HIF1α | late | 2 |
| Zhang et al (Zhang et al., 2010) | Body weight and AT mass ⇑ | aP2-Cre in C57BL6 mice | Dominant negative HIF1 | early-late | 3 |
| Sun et al (Sun et al., 2013) | Body weight and AT mass ⇓ | Adiponectin-Cre-rtTA | Dominant negative HIF1 | late | 4 |
| **HIF1 upregulation** | | | | | |
| Michailidou et al (Michailidou et al., 2015) | Body weight and AT mass ⇑ | ap2-Cre in C57BL6 mice | ap2-PHD KO | late | 5 |
| Drareni et al (Drareni et al 2018) | Hypertrophy | Adiponectin-Cre in C57BL6 mice | GPS2 KO | late | 5 |
| Halberg et al (Halberg et al., 2009) | Body weight and AT mass ⇑ | aP2-Cre in FVB mice | stblHIF expression | late | 6 |

| | | | | | |
|---|---|---|---|---|---|
| Rahtu-Korpela et al (Rahtu-Korpela et al., 2014) | Body weight and AT mass ⇓ | C57BL6 mice | PHD2 KO | early | 7 |
| Weiszenstein et al (Weiszenstein et al., 2016) | Lipids ⇑ | 3T3-L1 | 5% O2 | early | 8 |
| | Lipids ⇓ | 3T3-L1 | 1% O2 | early | 9 |
| Musutova et al (Musutova et al., 2020) | Lipids ⇓ | 3T3-L1 | 1% O2 (5 min cycles) | early | 9 |
| Sahai et al (Sahai et al., 1994) | Lipids ⇓ | 3T3-L1 | 3% O2 | early | 9 |
| Floyd et al (Floyd et al., 2007) | Lipids ⇓ | 3T3-L1 | PHD inhibitors | early | 10 |

**Related studies (not mapped)**

| | | | | |
|---|---|---|---|---|
| Thomas et al (Thomas et al., 2016) | Body weight ⇓<br>AT mass ⇑ | | PHD1 KO | early |
| Matsuura et al (Matsuura et al., 2013) | Body weight ⇓<br>AT mass ⇓ | ap2-Cre | ap2-PHD2 KO | late |
